# scSPARKL: Apache Spark based parallel analytical framework for the downstream analysis of scRNA-seq data

**DOI:** 10.1101/2023.04.07.536003

**Authors:** Asif Adil, Namrata Bhattacharya, Mohammed Asger

## Abstract

As the field of single-cell genomics continues to develop, the generation of large-scale scRNA-seq datasets has become more prevalent. While these datasets offer tremendous potential for shedding light on the complex biology of individual cells, the sheer volume of data presents significant challenges for management and analysis. To address these challenges, a new discipline, known as “big single-cell data science,” has emerged. Within this field, a variety of computational tools have been developed to facilitate the processing and interpretation of scRNA-seq data. In this paper, we present a novel parallel analytical framework, scSPARKL, that leverages the power of Apache Spark to enable the efficient analysis of single-cell transcriptomic data. Our methodology incorporates six key operations for dealing with single-cell Big Data, including data reshaping, data preprocessing, cell/gene filtering, data normalization, dimensionality reduction, and clustering. By utilizing Spark’s unlimited scalability, fault tolerance, and parallelism, scSPARKL enables researchers to rapidly and accurately analyze scRNA-seq datasets of any size. We demonstrate the utility of our framework through a series of experiments on simulated and real-world scRNA-seq data. Overall, our results suggest that scSPARKL represents a powerful and flexible tool for the analysis of single-cell transcriptomic data, with broad applications across the fields of biology and medicine.

## Introduction

Single-cell transcriptomic sequencing (scRNA-seq) has revolutionized the scientific understanding of the biological processes in an individual cell. Earlier, the cells in a tissue were considered to be homogeneous, however, scRNA-seq has proved otherwise. Cells at individual level exhibit heterogeneity, which leads to the various phenotypical variations in the overall tissue (Wu et al., 2017). Another advantage of scRNA-seq, is the identification of rare cells in the same tissues on the basis of various cell features which were unknown earlier.

With the advancement of the cell profiling techniques and rapid cost drops, scientists are now able to sequence millions of cells in parallel. 10x chromium profiling, for instance, provides the capability of profiling more than 80000 cells in one experiment parallelly(Bhattacharya et al., 2021). For each cell measurements of more than 20,000 genes are taken, generating a *Cell* x *gene* matrix of an enormous size. Illumina HiSeq, another important next-generation sequencing platform, can alone generate 100 GBs of raw transcriptomics data (Adil et al., 2021). It is expected that the genomic data may potentially surpass the data generated by social media giants like Facebook, Twitter (Adil et al., 2021). The generation of massive single-cell data at the rapid pace and on various experiment levels make the single-cell data science problem as a Big Data problem. This data boom needs to be managed, so that the analysis is fast and accurate.

The analysis of scRNA-seq starts from quality check of reads, alignment of reads and quantification of reads (generation of expression matrix); post quantification, we perform preprocessing on the gene expression matrix i.e., removing low-quality cell and gene filtering, normalization, selecting highly variable genes, dimensionality reduction, clustering and differential gene expression analysis etc. The processes from read quality check to generating an expression matrix of scRNA-seq is usually done with the help classic bulk RNA-seq tools. However, the remaining processes or the downstream analysis of scRNA-seq requires newly designed tools and algorithms.

Additionally, the atlas projects like Human Cell Atlas (Regev et al., 2017) are generating data in terabytes, hence increasing the demand of highly scalable and parallel architectural tools to handle the big data. The rescue can be found in big data analytics tools. The big data processing framework like Apache Spark and Hadoop are capable of handling terabytes of data efficiently (Salloum et al., 2016). To overcome the limitation of computing power for handling big single-cell data, we introduce a novel parallel computational pipeline backed by Apache Spark – a big data handling framework. The pipeline is scalable, extremely parallel and fault tolerant that runs on the commodity hardware.

### Apache Spark

Apache Spark is a distributed analytical engine made for handling big data. It provides an essential parallel processing platform for large datasets (Hildebrandt et al., 2020). Spark at its core uses an abstraction called Resilient Distributed Datasets (RDD), a read-only collection of objects distributed across the cluster of machines, which can be recreated if one of the machines is lost – fault tolerance (Salloum et al., 2016). All the jobs are launched in parallel through a program called “driver program” which also maintains the high-level flow control of the applications and jobs submitted to the executors. Spark uses in-memory concept i.e., it leverages Random Access Memory (RAM) of a machine and brings the data to be processed directly to the RAM thereby increasing the response time by removing the time taken to fetch data from a disc drive (Aziz et al., 2019). Since, RAM is faster than HDD it speeds up the execution of jobs. In the recent release of Apache Spark 3.0, GPU acceleration is facilitated to boost up the processing speed. Apache Spark applications can also be accelerated using the recently developed NVIDIAs RAPIDS-AI (Hricik et al., 2020). If GPU facilities are available, RAPIDS-AI provides a plugin for accelerating the Spark applications without changing the code.

## Materials & Methods

### Description of the datasets used

For testing the usability of the pipeline, we used the following datasets for experimentation:

#### Brain Non-myeloid Data

Tabula Muris senis, is a collection of single-cell transcriptomic data from the 20 tissues of Mus Musculus, with around 100,000 cells (Schaum et al., 2018). We used the scRNA-seq data of mouse brain non-myeloid cells. The data contains the expression measurements of 3401 brain cells for 23433 genes. The metadata file has the annotations for the dataset like sub-tissues: cortex, hippocampus, cerebellum and striatum; cell ontology class, mouse-id, and mouse-gender.

#### Jurkat-293T Cell data

The Zheng et al. data contains 3388 cells, from two species viz. mouse and human. The expression measurements of Jurkat and 293T cells, have been blended in vitro in equivalent proportion (50:50). Cells expressing CD3D are assigned Jurkat, whereas those expressing XIST are assigned 293T.

#### 68K PBMC

For time comparison across different data sizes, we used 68K Peripheral Blood Mononuclear Cells (PBMC) data from (Zheng et al., 2017). The cells have been collected from a healthy donor.

### Experimental Setup

The tasks were performed on Windows 11 OS, with 8GB RAM, intel i7 3.8Ghz processor with 8 cores. For development, we used Apache’s Spark version 3.1.2, Python 3.9 and JDK version 8.0. Table 1 (See Supplementary) provides the details of the hardware used in the experimentation. In Single-node Spark, one machine was used as a worker and the other one as master machine. In 2-node Spark environment, two machines were used as workers and one machine acted as a Master node. The memory allocation was done statically where each executor was provided with 2 GBs of memory. The rest of the memory is reserved for OS and Hadoop daemons.

### Preprocessing

We developed two exclusive Spark packages for loading data and data filtering. The data_load.py package file loads the data as a Spark dataframe and performs the initial preprocessing like removal of special characters such as periods (.), commas (,), blank spaces etc., from the columns and replaces them with the underscore (‘_’). The scRNA-seq data is usually in wide format and Spark has the ability to work better with tall/long format data; hence for optimization, we reshape the data to the tall format using *melt_Spark()* function in the pipeline. This step is essential for the users with low memory machines (i.e., as lows as 8GB). The dataframe is then written as a Spark Parquet file. Apache Parquet format is an open-source columnar storage file format which is based on “shredding and assembly algorithm”. It is designed to handle large and complex datasets for faster aggregation functions. The parquet formatted file is compressed, which speeds up the further analytical procedures on the file. We mainly use Apache Sparks aggregate functions, known for their quick retrieval and less computationally exhaustive/expensive. Pseudocode-1 provides the detailed outline of the data cleaning and reshaping.

#### Pseudo Code 1 Stage-1

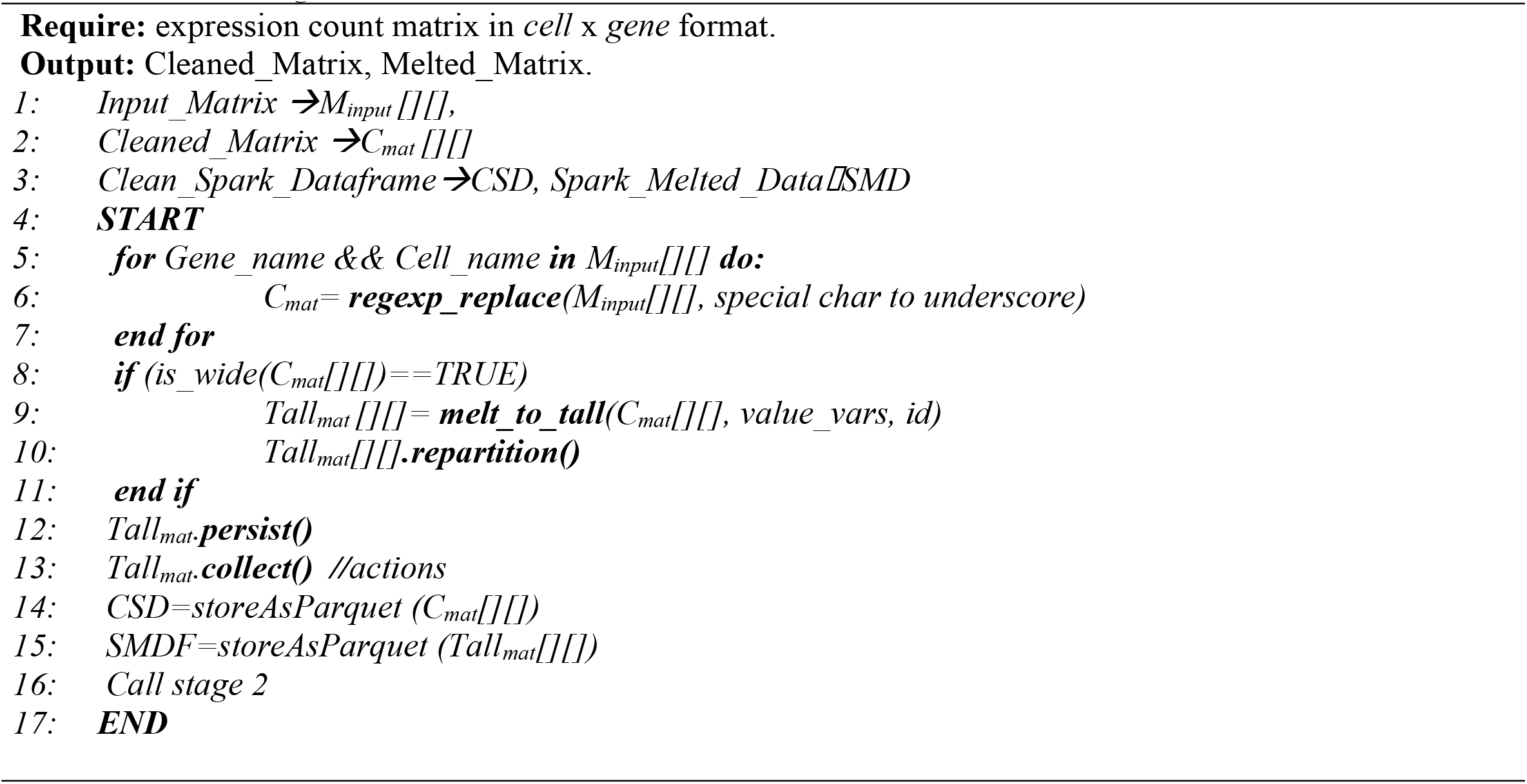

### Quality control: Gene and Cell filtering

For the coherent and accurate downstream analysis of scRNA-seq data, it is necessary to filter out low-quality cells and genes. We followed “scater” package (McCarthy et al., 2017), SCANPY (Wolf et al., 2018) and Seurat (Satija et al., 2015) for generating variety of quality control matrices. Quality matrix is calculated on number of parameters like total genes expressed per cell, percentage of ERCC, number cells expressing genes greater than zero, mitochondrial percentage detected in each cell etc., (Supplementary Table 1 & 2). We calculate the log measurements and mean measurements of numerous parameters for the easier interpretation such that a good parameter for cell and gene filtering is defined. In addition to the normal filtering procedures, we have also implemented a Spark based Median Absolute Deviation (MAD) filtering methodology (see supplementary) for filtering out the genes with a normalized count value greater than a decided threshold (usually >3). Pseudocode-2 provides an outline for the whole process of cell and gene quality measurements, and pseudocode −3 outlines the filtering process on the basis of cell and gene quality summaries. Further, pseudocode-3 also includes the process of normalization, dimensionality reduction and clustering, which are described in following sections.

#### Pseudo Code 2 Stage-2

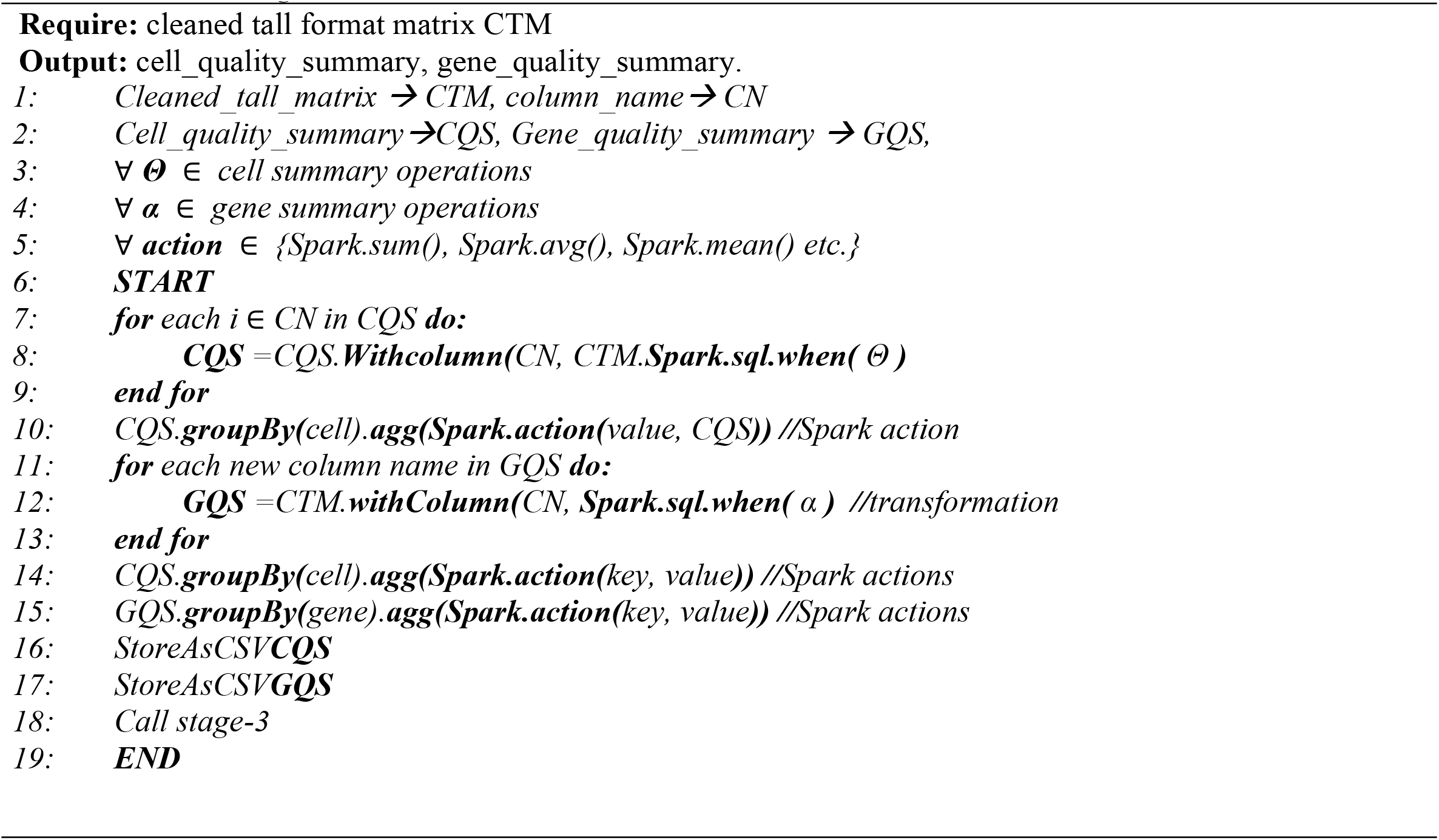

### Normalization

The scRNA-seq data is highly sparse, it is necessary to normalize and scale this data so that the biological inferences are accurate and further downstream analysis are crisp. The *data_normalize*.*py* file of the framework currently implements two simple and straightforward normalization methods: Quantile normalization (QN) and Global Normalization (GN).

In QN, the genes across each sample/cell are ranked, lowest to highest, as per the expression measurement and the resultant ranks are retained. The dataframe is arranged according to the ranks of the rows. These rows are then averaged and an average matrix is prepared. The earlier retained rank matrix is used to reposition the obtained averages at their respective place. This makes it easier to compare the values of different distribution, while preserving the actual coherence of the matrix.

The Global normalization is similar to converting to CPM. In this, we divide each expression value of a cell by the total number of counts of that gene. The resultant value is then multiplied by 10,000 and then log transformed with the addition of a pseudo-count 1. Mathematically, if *X*_*m*n*_ is an unnormalized filtered scRNA-seq matrix, then the Normalized matrix *X*_*N*_ can be given as:

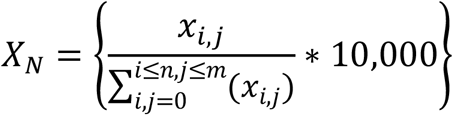

*where x* ∈ *X*_*m*n*_ *and 10,000 is a scaling factor* and this scaled normalized matrix is then log transformed as:

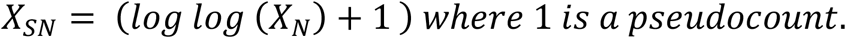

### Dimensionality Reduction

Dimensionality reduction is the process of projecting primary features of a dataset, from higher to lower dimensions, while trying to preserve the originality of the data (Ayesha et al., 2020). scRNA-seq data, falling in with the criterion of Big Data, gets affected by the curse of dimensionality (Feng et al., 2020). It is impossible to study the measurements in higher dimension dataset. For dimension reduction, we use Principal Component Analysis (PCA), t-distributed Stochastic Neighborhood Embeddings (t-sne) and Uniform Manifold Approximations and Projections (UMAP). Currently, we implement the python-based versions of UMAP and t-sne which partially work independently from Apache Spark; however, they use Apache parquet as a storage technology. The PCA, however, is exclusively Spark based and is equally parallelizable across the distributed units.

### Clustering and further downstream analysis

One of the important advantages of scRNA-seq is the identification of unknown and known cell clusters (Menon, 2018). The single-cell profiling techniques have undergone a variety of advancements, leading to the improvement in capture efficiency, which in-turn has resulted in sequencing of cells on a multi-million scale and hence a multi-million scale dataset. This demands the better computational requirements for the unsupervised clustering of these cells (Baker et al., 2021).

The K-means clustering intends to group the given *n* observations into *k* clusters (*k<=n)* having minimum within-cluster variance. It divides the input datapoints into k distinct clusters iteratively, such that these data points converge to a local minimum. As a consequence, the clusters created are compact and self-contained. It works in two stages 1) selecting the *k* centroids randomly, 2) and bringing the datapoints to the nearest possible center. The *k* in k-means is chosen using elbow plot method or by calculating silhouette score, and both are provided for the user in the framework.

The next step after clustering is the Differential Gene Expression (DGE) analysis. We use two-sample T-test to determine top 10 significant genes in the obtained clusters, which are ranked based on their p-value (<0.05).

#### Pseudo Code 3 Stage-3

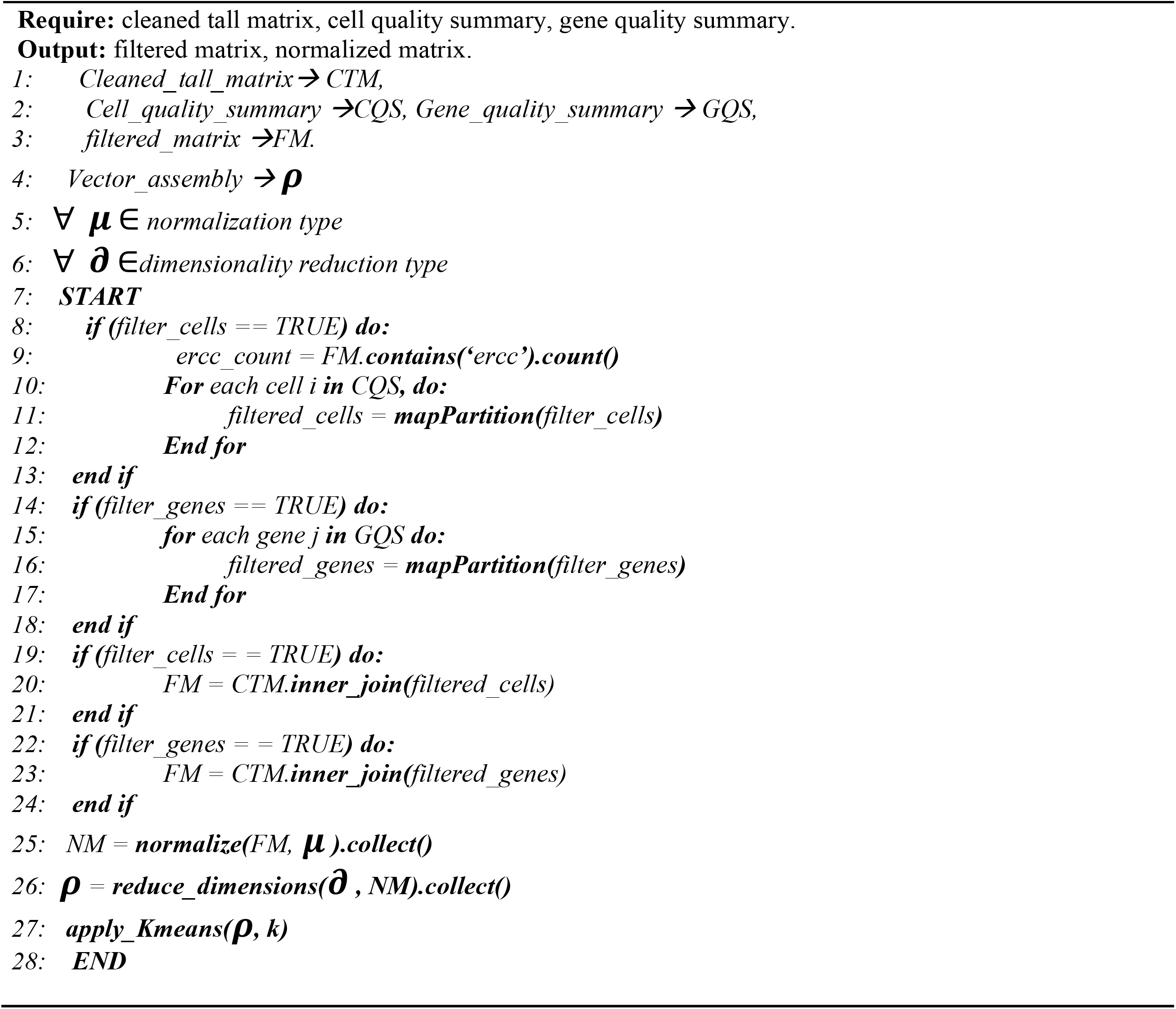

## Results

The scSPARKL pipeline is extremely fast and accurate. It is the first Apache Spark based computational and analytical framework for scRNA-seq data. With apache Spark acting as a core engine, enabling the parallel utilization of the resources, it takes less than 10 minutes to perform filtering, normalize, dimension reduction, clustering and visualization of the data. The reanalysis results produced by our framework are similar to what are presented in the original authors (Schaum et al., 2018) and (Zheng et al., 2017) with total computation time elapsed as low as 8 minutes. The mouse brain cells, analysis shows the abundance of oligodendrocyte and endothelial cell type in brain non-myeloid cells, as has been found by the original authors (supplementary S2, S3, S5, S6). The framework source-code is fully open source and is available to the community for further necessary modifications and extensions.

### Scalability and performance improvement with Apache Spark

The data for performance evaluation was generated from PBMC. We divided the data into five randomly sampled sets of 2000, 4000, 6000, 8000, and 10,000 cells and each dataset has more than 30,000 genes. This was done to measure the time consumption and scalability of the pipeline across different cell loads.

For comparison, we setup a sequential/linearly working python-based standalone pipeline (SSA) having similar functionalities. The comparison was drawn between an SSA, single-node Spark environment, and 2-node Spark environment (Supplementary Table 3). With the increase in nodes, As can be seen in the figure 3, the largest (10K) chunk of data takes ∼10 & ∼13 minutes to produce the final output in a 2-Node Spark cluster and Single-Node Spark environment, respectively. While as the same dataset takes more than 50 minutes, to produce the final output, on a non-Spark SSA. This indicates the performance improvement of 80% and 74% using 2-node and single-node Spark clusters, respectively, for all datasets. We anticipate that with an increase in physical cores and good RAM, the speed up can go beyond 100x.

**Figure 1:**
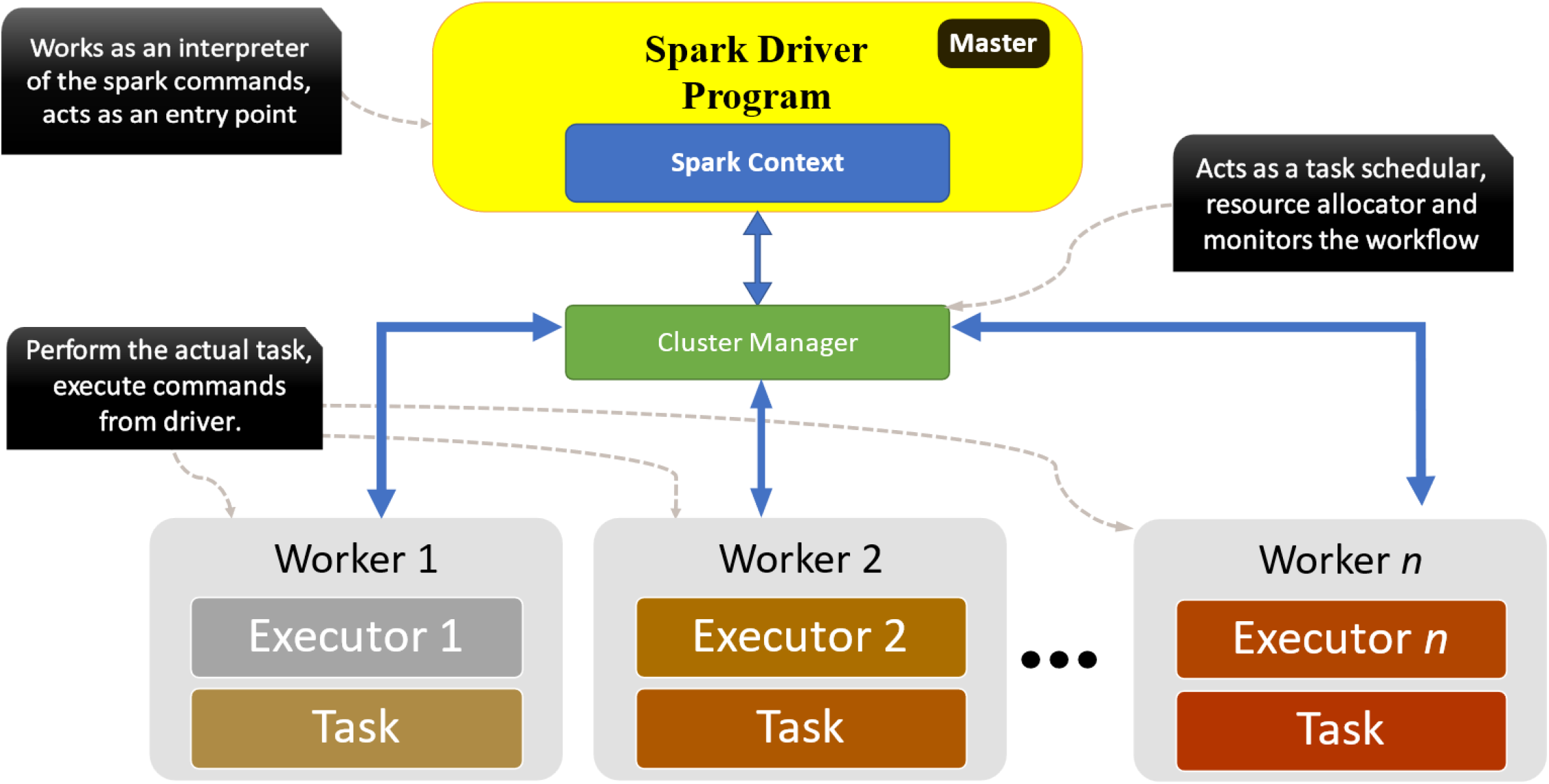
Apache Spark workflow: the Spark driver program works as a master and as an entry point for all the Spark jobs. The master submits jobs to the worker nodes. The cluster manager keeps the track of the nodes and the jobs distributed to them, several cluster managers are Yet Another Resource Negotiator (YARN), Kubernettes, mesos and standalone (in our case). The worker/slave nodes are the actual machines where the tasks are executed and they report back to the cluster manager

**Figure 2:**
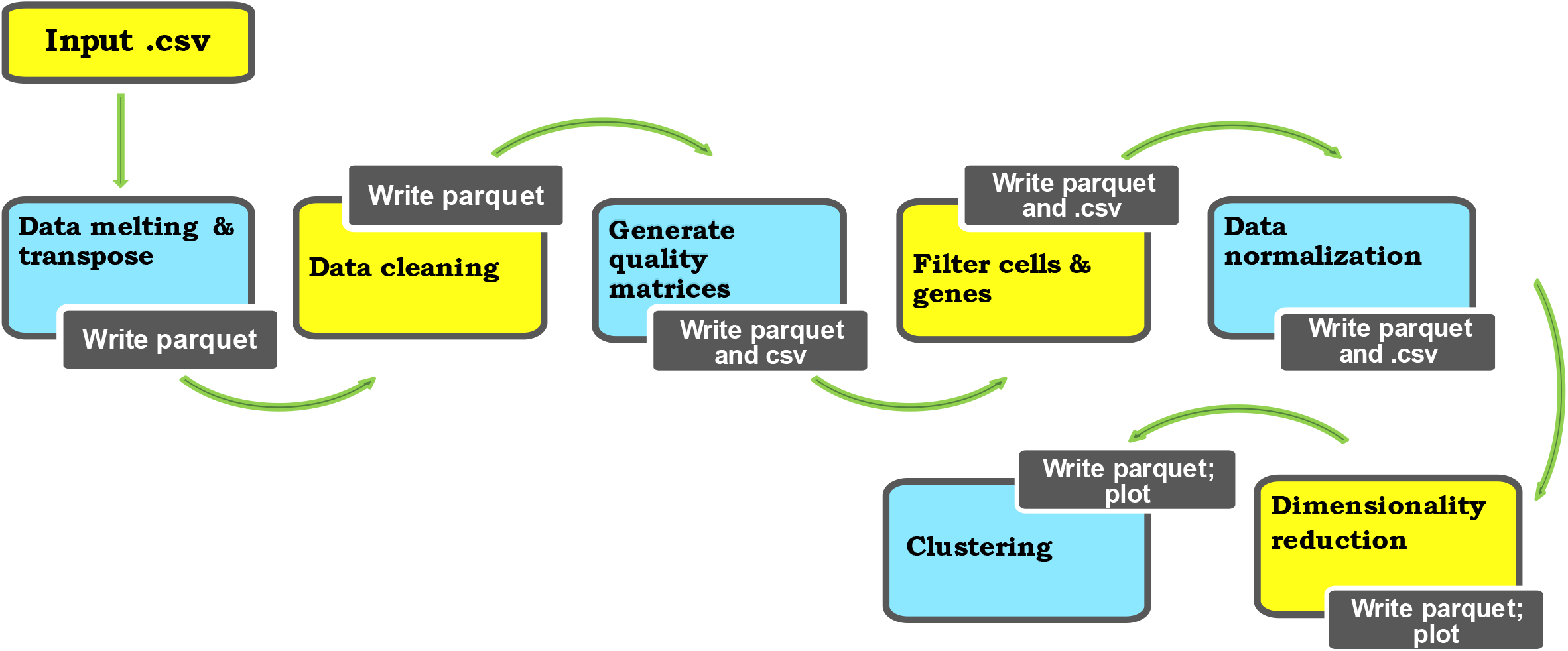
General workflow of the scSPARKL pipeline. Described with full details in materials and methods.

**Figure 3:**
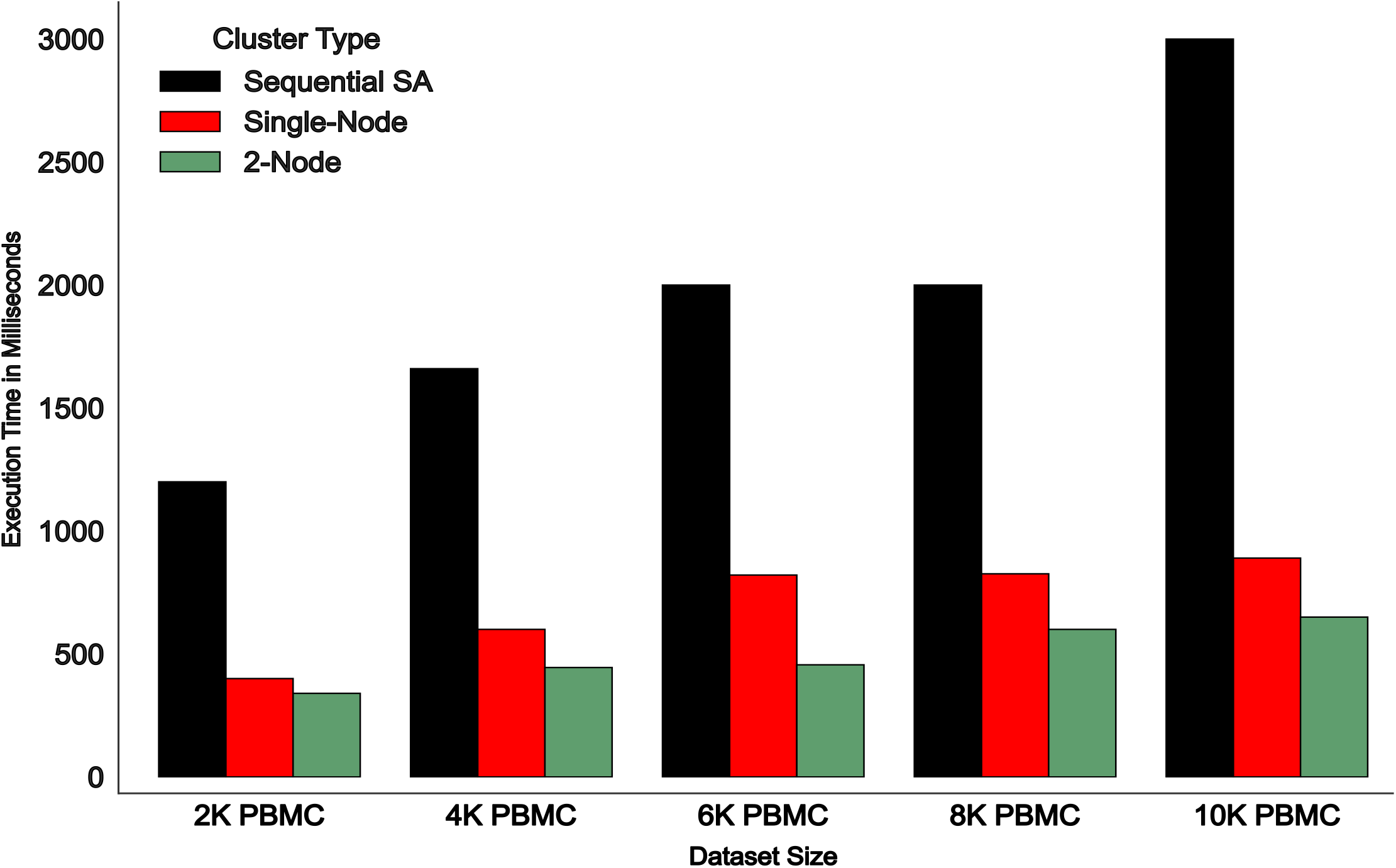
Bar Plot representing the total time taken (in seconds) from submitting the job to the final output i.e., clustering, for different data with different cell loads. Cells were randomly sampled and chunked from 68K-PBMC data.

We observed (Supplementary figure S7) that persisting a dataframe improves the execution time of the processes like generating quality summaries and filtering the low-quality cell and genes, this however occupies much of the RAM of an executor. We also observed, the usage of parquet as a data storage format was also a good choice, since it reduces the shuffle-read time.

Although the Spark foundations were laid on RDDs, the java serialization of jobs and garbage collection raise the memory overhead and prove to be disadvantageous in case of low memory. We, instead of using RDD, exploited Spark Dataframe API and used SQL functions for the number of operations, thereby improving much of the execution time.

### scSPARKL conforms with original methods

For validation of results, we performed the reanalysis of Brain Non-myeloid cells and Jurkat-293T Cell data, using scSPARKL. The analysis produced similar results as produced by the original authors, however with in lesser time (figure 4).

**Figure 4:**
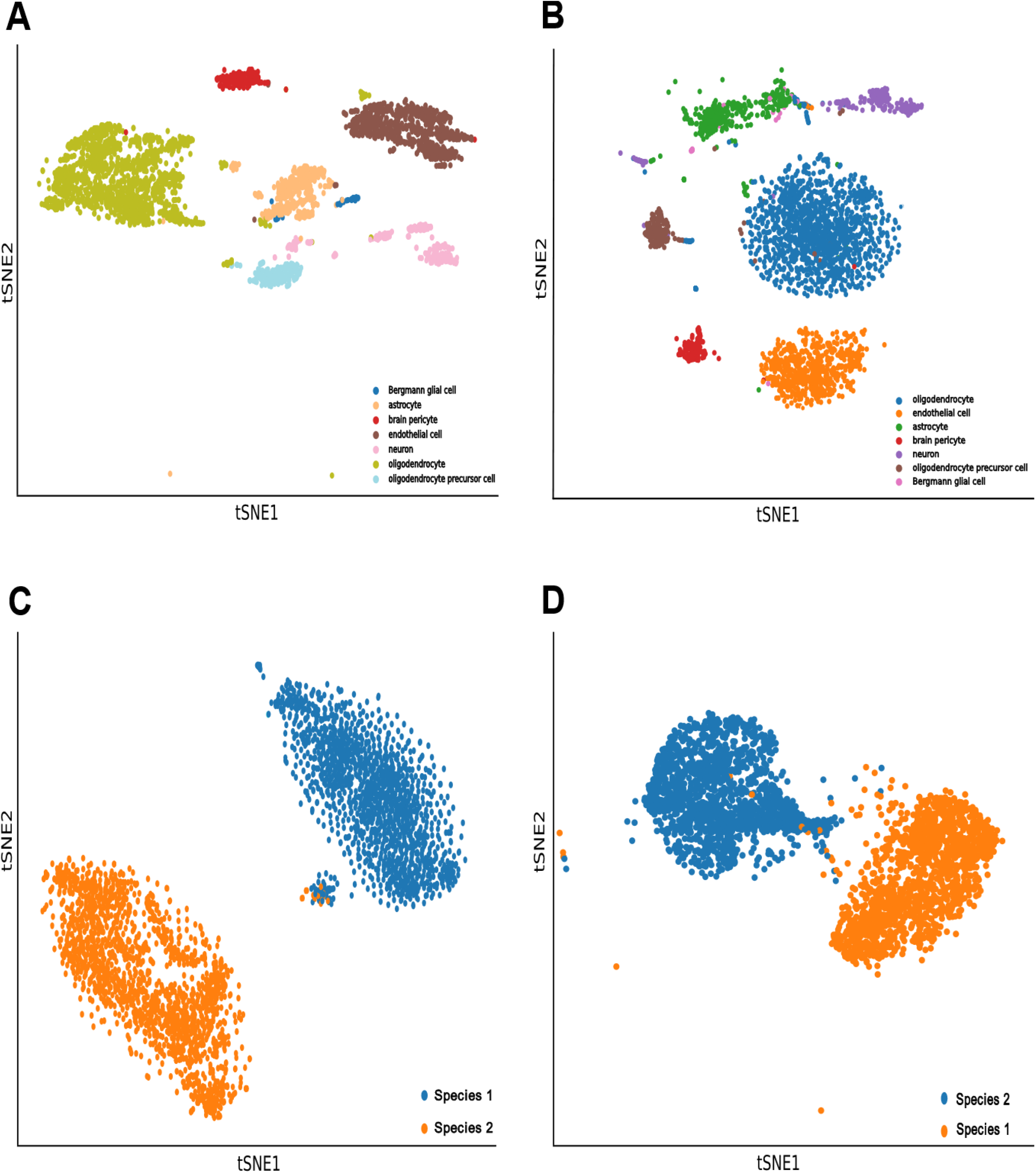
**A)** tSNE visualization for Brain Non-Myeloid data from SCANPY. **B)** tSNE visualization of Brain Non-Myeloid cells using scSPARKL. **C)** tSNE visualization for Jurkat Cell data from original author. **D)** tSNE visualization of Jurkat-239T Cells using scSPARKL.

As can be seen in the figure 4b and 5b, the framework is able to differentiate the cell types much accurately. The outliers in the plots can be attributed to the less strict filtering of cells and genes and different parameters for tSNE algorithm. The plots, from the proposed framework are with default filter parameters for cells and genes removal these are: ‘percent_ercc > 10%’, ‘percent_mito > 5%’, ‘cells_expressing < 10 genes’, ‘genes expressing in < 3 cells’, ‘dropout rate > 70%’.

### Adjusted Rand Index for cluster performance evaluation

The Adjusted Rand Score (ARI) for the clusters is used to establish a comparison between clustering results. The ARI for Jurkat-293T cell line and Brain non-myeloid cells using scSPARKL is ∼0.90 and ∼0.50 respectively (Figure 5). The value of *k* was set as per the number of cell types in the data.

**Figure 5:**
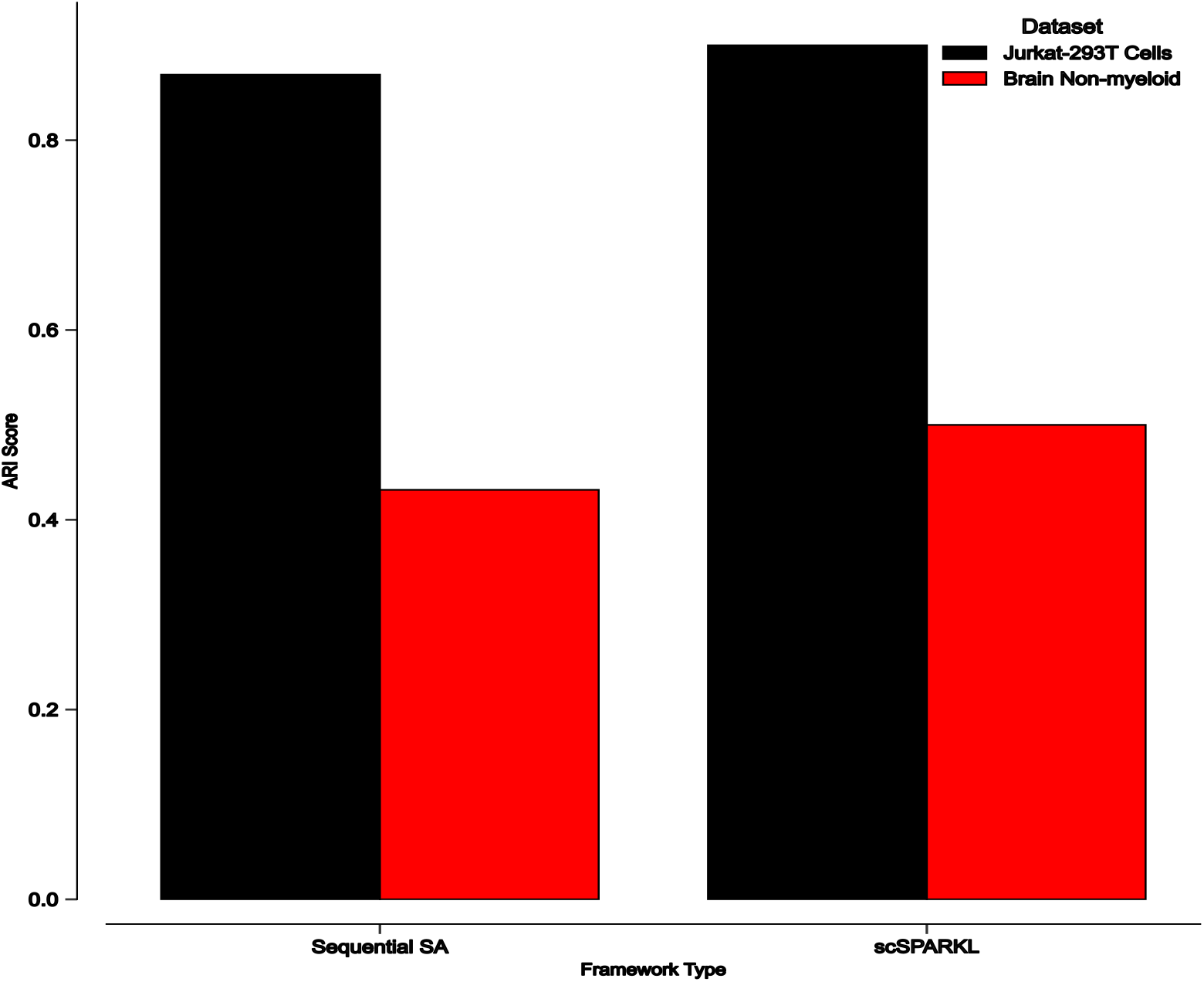
ARI scores of Clustering results from Sequential Standalone scRNA pipeline and scSPARKL

## Discussion

Single-cell transcriptomic data is increasing phenomenally, the data accumulation from scRNA-seq demands efficient tools to handle the data explosion. Since single-cell data comply to Big Data characteristics, we developed an Apache Spark based analytical pipeline for the handling of scRNA-seq data. Our pipeline requires a .csv file as an input. The input is checked for the tall and wide format, as Spark works better with tall format data, the input is melted from wide format to tall. This data is stored as a parquet file for further perusal. This is followed by the data cleaning, i.e., removal of incomprehensible characters from the attributes of data like commas, parenthesis, dots etc., if they are present. For differentiating the names, we replace all these special characters with underscore.

After cleaning, the next step is preprocessing i.e., we generate a quality check dataframes, based on the various quality parameters, like total amount of genes expressed for a cell, percentage of ERCC’s and percentage of mitochondrial genes expressed. Similarly, a gene quality dataframe is generated for the data. These dataframes are stored separately, and a parquet file for the two is generated again. This makes it easier for the user to take up analytical steps from the middle, avoiding further data melting. Based on the quality check matrices a user can decide a threshold for cell and gene filtering, depending upon various factors, and the cell and gene filtering is done by calling the *data_filter*. This is again saved as a parquet file and a separate .csv file is also generated for future usage.

After filtering cells and genes, we have developed a parallel version of normalization. We have currently kept two most common and widely used normalization methods for a user to utilize, these are quantile normalization and global normalization (also called as simple normalization). The gene selection can be done by either calling the top n highly variable genes by calculating their coefficient of variance or by calculating the Median absolute deviation with a deviation of greater than 3.

Single-cell suffers from curse of dimensionality and it is necessary to reduce the dimensions to have a holistic view of the data. The dimensionality reduction, in our framework, can be performed using a parallel version of PCA. We also provide a pySpark based versions of UMAP and t-sne for the dimension reduction. The choice remains with the user depending upon the availability of computational power. Using UMAP for dimensionality reduction ensures that the global structure of the data is preserved and it is relatively faster than t-sne (Adil et al., 2021).

Implementation of Spark as a core engine ensures unbounded parallelism, fault tolerance and fast computation of unlimited data. Other established tools like scanpy uses Pandas as a base package for data handling, however, pandas cannot handle terabytes of data and cannot be parallelized. This makes our tool a novel amongst all others.

As Spark is based on in-memory computation, a limitation of our framework can be the requirement of higher amounts of RAM to work in a standalone environment on a single computer.

For now, the limitation is managed using data melting –a procedure to combine a group of columns from a wide formatted data into a single column of data. However, the computational steps after normalization require wide format data, and this wideness may reach more than 20000 columns, which increases the requirement for a higher memory. This can be resolved by running the Spark in cluster mode with each node having a separate memory, the Spark jobs tend to run faster with higher degree of parallelism.

## Conclusion

In light of the unprecedented accumulation of scRNA-seq data over the past few years, we have developed a parallel computational framework that offers an intuitive and user-friendly approach to data management and subsequent analysis procedures. Our framework is a viable alternative to existing tools that lack scalability and fault tolerance, since the majority rely on pandas, which can collapse and stutter as data size exceeds 50GBs. By utilising Spark at its foundation, we ensure that our framework can process terabytes of data at a breakneck pace. With the advent of single-cell studies at the proteomic and genomic levels (e.g., CITE-seq, scATAC-seq, and scBS-seq for single-cell methylation), we plan to develop Spark-based pipelines for the integration of single-cell multi-omics data. Our framework represents a major advancement in the analysis of single-cell transcriptomic data, with broad applications in the disciplines of biology and medicine.

## Supporting information

(See Supplementary)

## Code Availability

The framework implementation and the source code, using PySpark, can be found on following: https://www.github.com/asif7adil/scSparkl

## Conflict of interest

None Declared.

## Acknowledgement

The authors are highly thankful to Prof. Debarka Sengupta (Computational Biology division of IIIT-D, India), for his valuable inputs and suggestions which improved the overall quality of the manuscript.

## Funding

No funding was availed for the study.

## Notes

### Competing Interest Statement

The authors have declared no competing interest.

