## Supplementary material for "scSPARKL: Apache Spark based parallel analytical framework for the downstream analysis of scRNA-seq data": (See Supplementary)

### Section 1: Cell and Gene quality matrix quality measures according to which the parameters for gene and cell filters can be set.

**Table 1:** Cell Quality measures

| S.No. | Cell quality measure |
|---|---|
| 1 | Sum of total expression per cell |
| 2 | Total number of cells with expression count > 0 |
| 3 | Average expression count in each cell |
| 4 | Total ERCC count present in cell |
| 5 | Sum of ercc |
| 6 | Mitochondrial entries for each cell |
| 7 | Sum of mitochondrial counts in a cell |
| 8 | Percentage of ercc measurement in each cell |
| 9 | Percentage of mitochondrial expression measurement |
| 10 | Log1p of sum of total expressions in cell |
| 11 | Log1p of Sum of Ercc measurements |
| 12 | Log1P of sum of Mitochondrial genes |

**Table 2:** Gene Quality Measures

| S.No | Gene Quality Measures |
|---|---|
| 1 | Number of cells where gene count is greater than 0 |
| 2 | Total Sum of expression counts in each cell |
| 3 | Total no of dropouts for a gene |
| 4 | Percentage of dropout by counts |
| 5 | log1p_of_mean_counts |
| 6 | log1p_total_counts |

### Section 2: Extended results.

### S2.1 Jurkat-239T Visualization:

The gene and cell quality summaries for Jurkat-239T data are given in the figure S1.

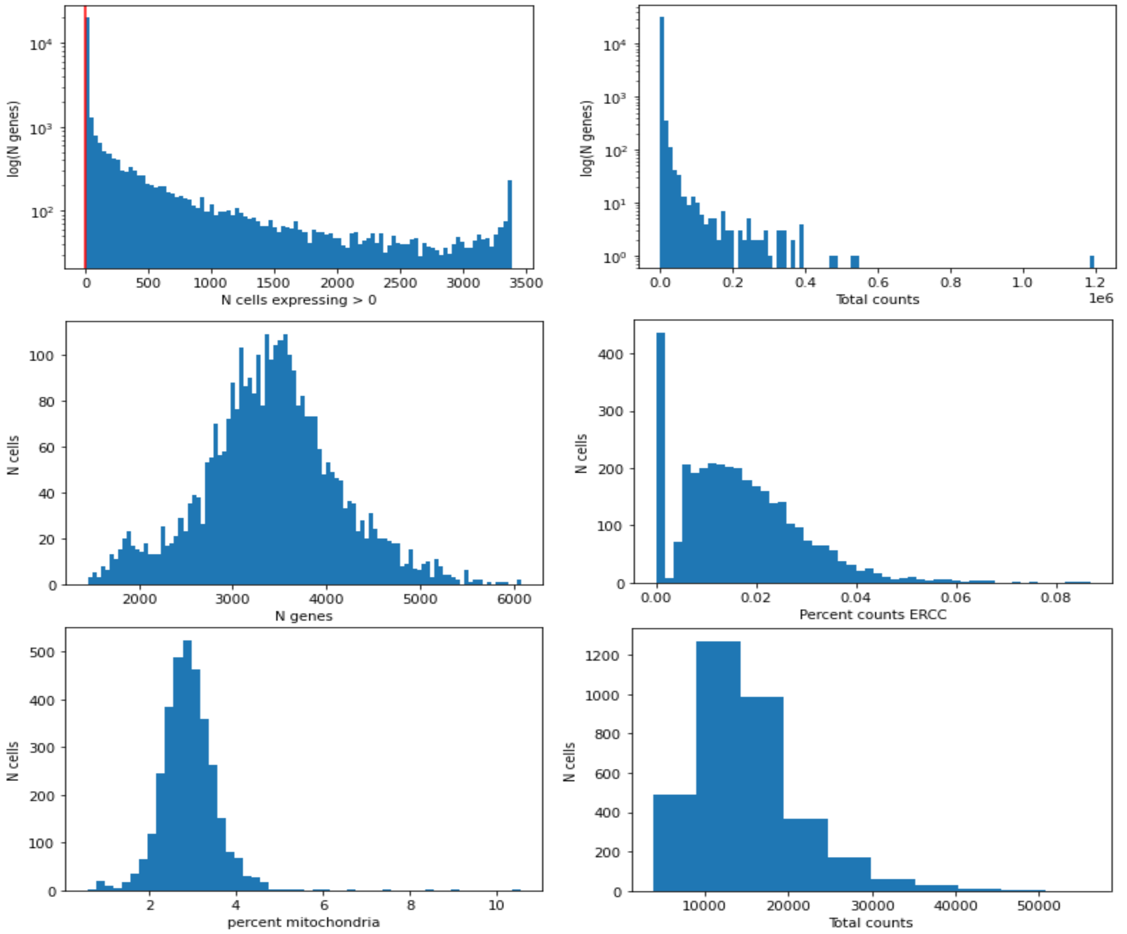


**Figure S1:** Histogram of Gene and cell qualities, respectively, for Jurkat-239T. This is used for deciding the filtering thresholds, depending upon the nature of data as well as the biological question in hand.

### S2.1.1 UMAP visualization

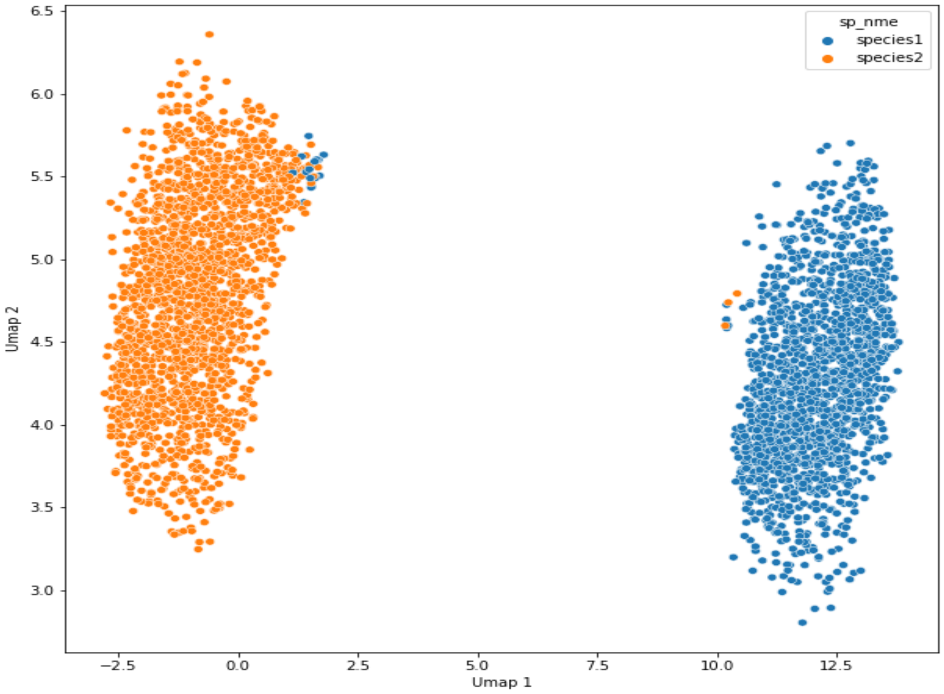


**Figure S2:** UMAP visualization of Jurkat-239T cells using scSPARKL with default filtering parameters.

### S2.1.2 Clustering Analysis for Jurkat-293T cells:

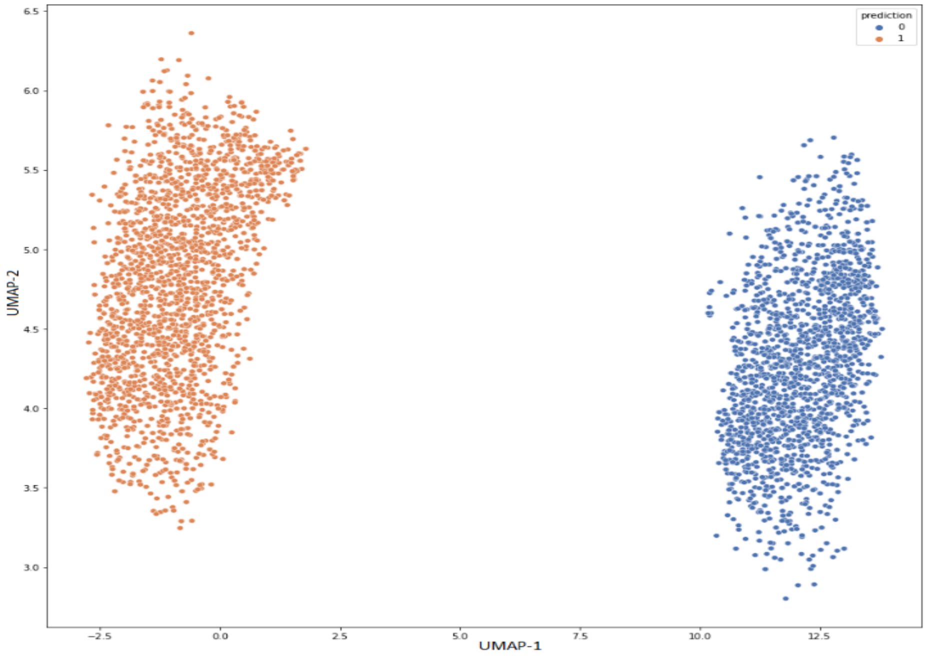


**Figure S3:** Clustering analysis using K-means clustering algorithm with k=2, silhouette score = 0.92145, ARI score=~90.

**S2.2 Mouse Brain Non-Myeloid Cells visualization:**

The gene and cell quality summaries for Mouse Brain Non-Myeloid data are given in the figure S3

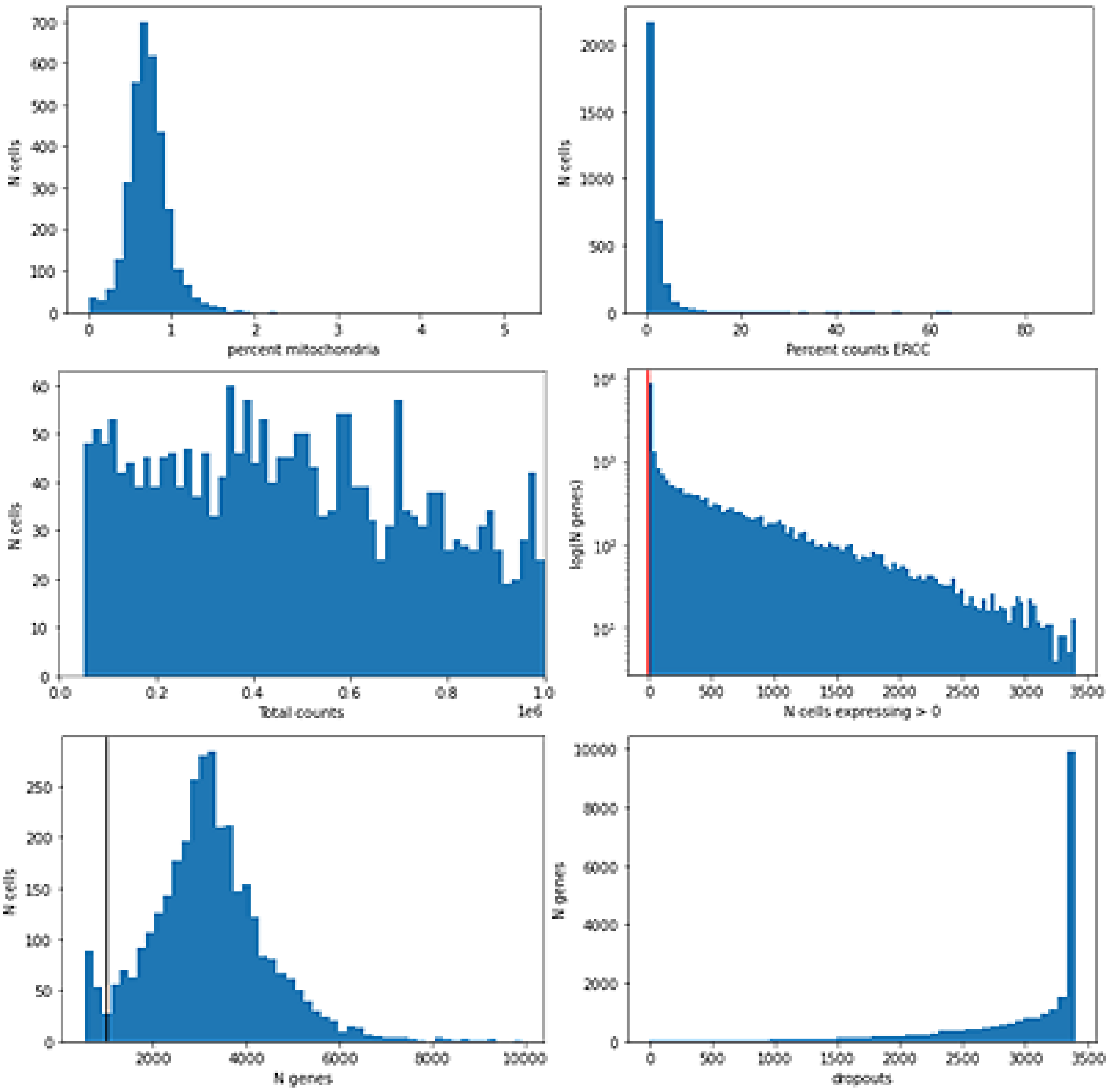


**Figure S4:** Histogram of Gene and cell qualities, respectively, for Mouse Brain cells. This is used for deciding the filtering thresholds, depending upon the nature of data as well as the biological question in hand.

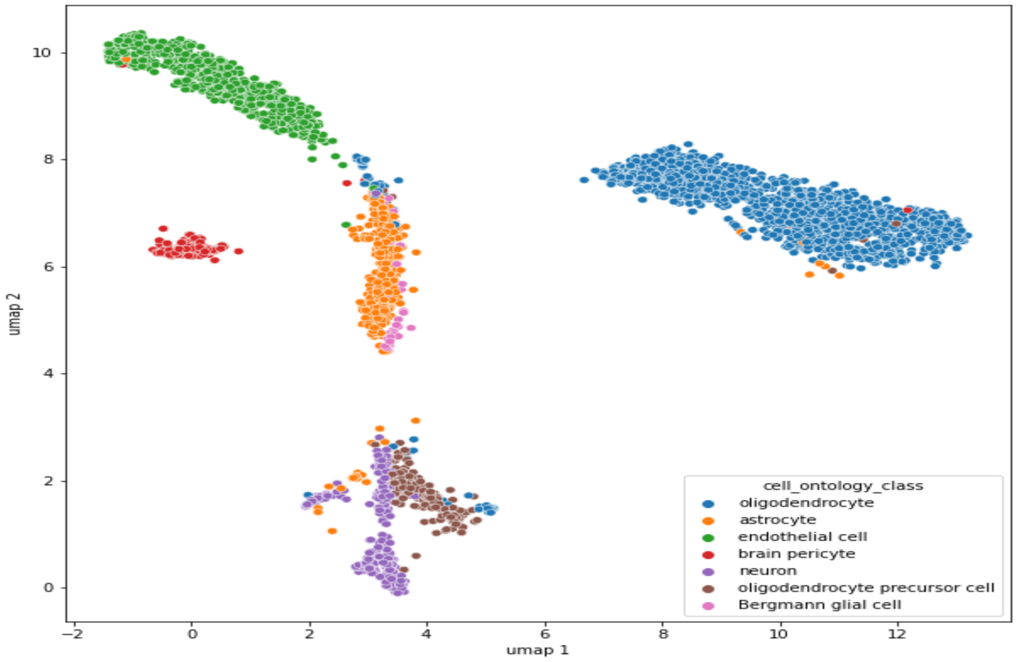


**Figure S5:** UMAP visualization of Mouse Brain Non-Myeloid data, with default filter values for cell and gene filtering.

### S2.3 Clustering of the Tabula Muris data.

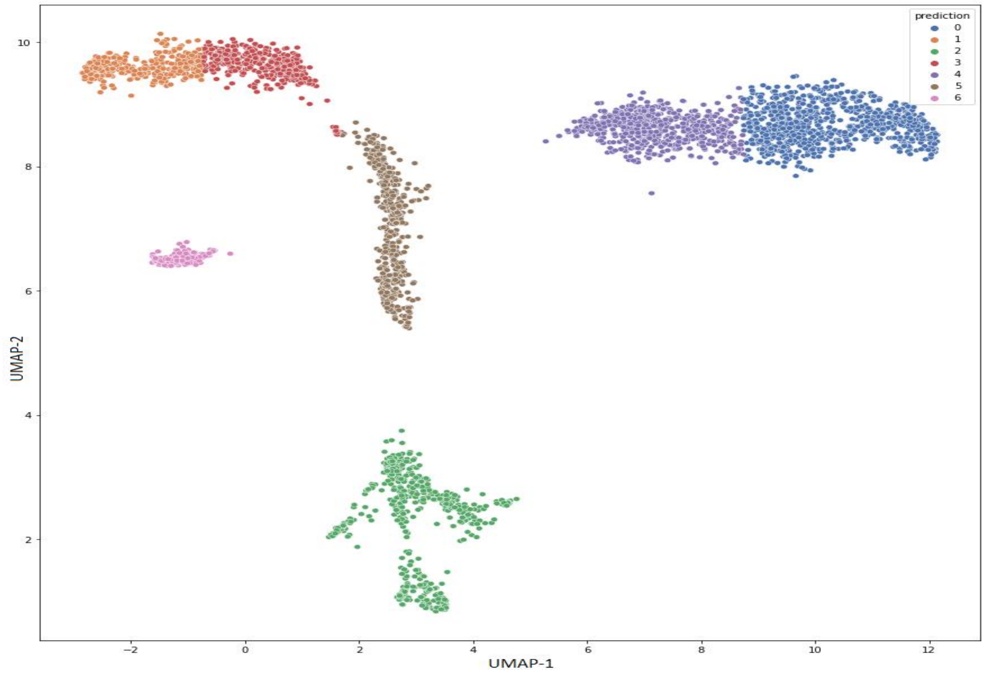


**Figure S6:** Clustering analysis based on UMAP embeddings of Mouse Brain Non-Myeloid cells, with highest silhouette score = 0.82952 and ARI score=~50.

### S2.3: Performance of individual processes:

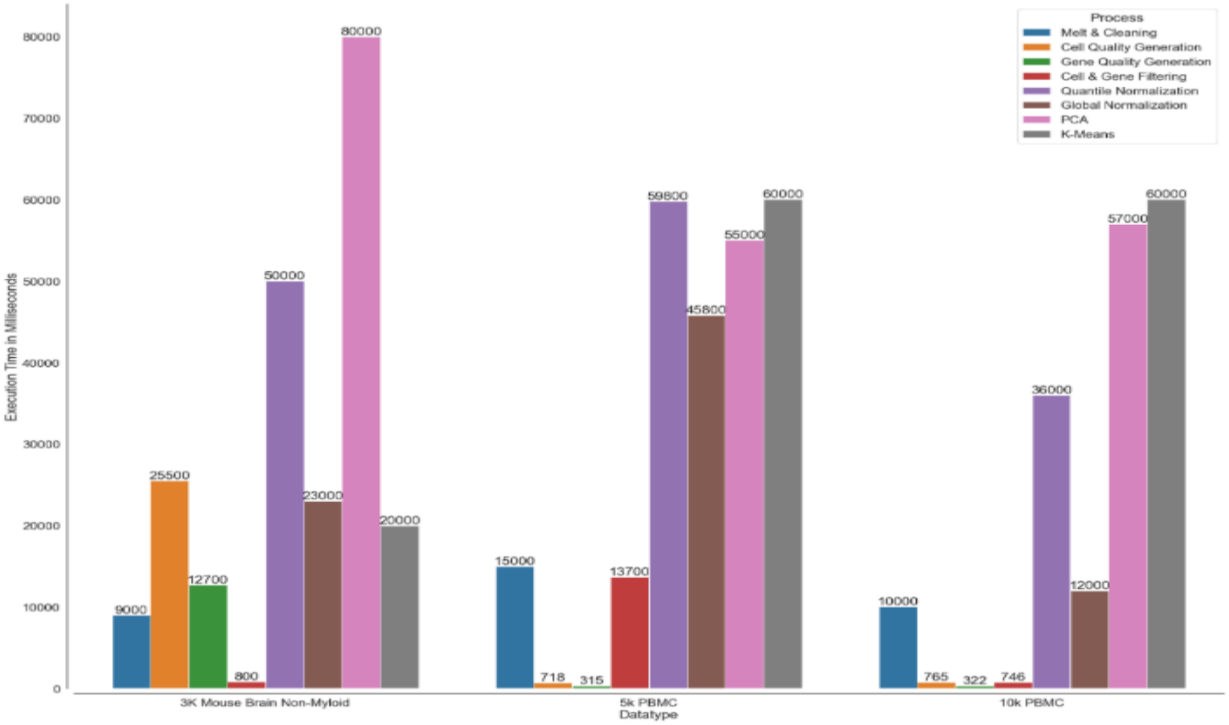


**Figure S7:** Bar Plot depicting the time (in Milliseconds) taken by each process on different data sizes. All the processes shown in the plot are exclusively Spark based, running parallelly on each core. UMAP and t-SNE are not shown here, since they are not Spark based. The processes are not inclusive of read/write operations. The features for PCA were 1200 for PBMC data, and 9100 for Mouse Brain data. Melting and Cleaning operations were measured as per the wall clock time.

### Section 3: Hardware information

**Table 3:** Details of the hardware used in the experiments.

| Environment | Master/Slave platform | Cumulative RAM | No. of Cores | Processor type |
|---|---|---|---|---|
| Python sequential | × | 6GiB | 4 | i7, 11th gen |
| Single-Node Spark | ✓ | 6GiB | 4 | i7, 6th gen |
| 2-Node Spark | ✓ | 6+6 = 12GiB | 8 | i7, 5th gen and 6th gen |
